# Contribution of larvae developing on corn and dry beans to the adult population of western bean cutworm in Michigan

**DOI:** 10.1101/2020.06.19.162016

**Authors:** Dakota C. Bunn, Eduardo Dias de Oliveira, Frederick Springborn, Miquel A. Gonzalez-Meler, Nicholas Miller

## Abstract

The western bean cutworm, *Striacosta albicosta* (Smith) (Lepidoptera: Noctuidae), is historically a pest of both corn (*Zea mays* L.) and dry beans (*Phaseolus* sp L.) in the western Great Plains. However, it has recently undergone an eastward range expansion establishing itself across the Corn Belt in twenty-five states and four Canadian provinces. To mitigate the effects of infestation in Michigan, foliar insecticides are used in dry beans whereas management of the pest in corn relies more heavily on the use of Bt-expressing hybrids. In this study stable carbon isotope analysis was used to determine what crop adult moths developed on as larvae with analysis showing that very few of the adult moths developed on dry beans. These results suggest that beans and corn are not suitable as co-refuges and that mainly adults which developed on corn are contributing to the next generation of western bean cutworm in Michigan.

## Introduction

The western bean cutworm, *Striacosta albicosta* (Smith) (Lepidoptera: Noctuidae), is a univoltine pest of both corn (*Zea mays* L.) and dry beans (*Phaseolus* sp L.). In corn, feeding with an infestation of one larva per ear can result in a loss of four bushels per acre, with large infestations reducing the yield by 30 to 40%, while in beans the yield is reduced by 8 to 10% (Peairs 2014). Additionally, feeding provides entry for molds and other plant pathogens, increasing the risk of diseases and mycotoxin contamination (Rice and Dorhout 2006, Seymour et al. 2010, Difonzo et al. 2015). Historically a pest of the western Great Plains region of the United States, over the past quarter of a century it has expanded eastward and is now established in at least twenty-five states and four Canadian provinces (Hoerner 1948, Hagen 1962, Jeschke 2018). In Michigan, the western bean cutworm was first detected in 2006 and severe injury to field corn was observed the following season (Smith et al. 2018).

While the western bean cutworm has greatly expanded its geographical range, little is known about its seasonal and local flight, however, cutworms are generally known to be strong fliers. For example, the army cutworm (*Euxoa auxiliaris* G.) migrates from Montana to Wyoming to reach summer aggregation sites while the black cutworm (*Agrotis ipsilon* H.) engages in long distance seasonal migrations to overwinter in warmer climates (Hendricks 1998, Showers 1997). However, unlike the army cutworm and the black cutworm, the western bean cutworm overwinters underground and the possibility of a seasonal migration of this pest has yet to be studied (Blickenstaff 1979, Smith et al. 2018).

Western bean cutworm prepupae overwinter before pupating in the spring. Adults then emerge and lay eggs from the end of June to mid-August (Blickenstaff 1979). In corn, oviposition occurs near the whorls on the upper side of the leaf just before tasseling. Once hatched, the larvae will feed on the tassels before migrating into the ear and feeding on the silks and kernels for the remainder of their development (Hagen 1962). In dry beans, eggs are laid on the underside of the leaf and once hatched the larvae feed nocturnally on the leaves and buds of the plants. Once further developed, the larvae feed on the developing seeds and pods at night while moving to the soil during the day. On both plants, once the larvae reach the sixth instar, they burrow into the soil and create overwintering chambers before the process starts anew (Hoerner 1948, Seymour et al. 2010).

While the western bean cutworm may feed on two very different plants it is not a generalist herbivore. Larvae have shown some success developing on several species of *Phaseolus* and some related legume species, but they cannot develop on soybeans (*Glycine max* L. Merr.) and often develop poorly on other cultivated legumes. Additionally, they cannot develop on the ancestor of domesticated corn, teosinte (*Zea mays* L. ssp *mexicana*) or other cultivated Poaceae. Finally, they are unable to develop on Solanaceae, including tomatoes (*Solanum lycopersicum* L.) (Blickenstaff and Jolley 1982). Thus, western bean cutworm appears to be unusual in being a specialist on two unrelated crop hosts.

Management of the western bean cutworm varies depending on the host and time of the growing season. In beans, foliar insecticide applications are the main mechanism of control, with pyrethroid applications showing excellent control of the western bean cutworm (Difonzo et al. 2015). However, application of foliar insecticides requires labor intensive scouting practices which are challenging due to the egg masses being located on the underside of the leaves and the nocturnal habits of the larvae. Another scouting method requires the use of pheromone traps to capture adult moths to determine when insecticide treatment is needed. Unfortunately, established insecticide application thresholds have been unreliable in the Great Lakes region where significant bean and corn damage are observed when trap numbers are low, compared to those observed in native western regions such as the state of Nebraska (Difonzo et al. 2015). In corn, foliar insecticide treatments are occasionally used, but they are ineffective once the larvae have entered the ear (Smith et al. 2019). Due to this, transgenic corn hybrids are a widely used approach to western bean cutworm management.

Transgenic corn hybrids expressing proteins derived from *Bacillus thuringiensis* are used to manage western bean cutworm infestation. Previous studies have demonstrated that Cry1Aa, Cry1Ab, Cry1Ac, and Cry9C proteins are unable to control WBC and that while Cry1F was once effective, WBC has gained tolerance to the toxin (Dyer et al. 2013, Ostrem et al. 2016, Wolff 2018, Smith et al. 2019). Therefore, the only current hybrids that are marketed for successful control of WBC are the ones expressing Vip3A (Difonzo et al. 2018, Farhan et al. 2018, Smith et al. 2018, 2019). Varying management strategies for the two crop types grown in near proximity to each other highlights a need for further development of insect resistance management (IRM) programs to mitigate the development of insecticide resistance. This subsequently requires further understanding of WBC biology and interactions between adult moths that developed on different host crops.

Currently IRM programs recommend the use of refuges in corn in order to sustain a population of susceptible target pests (Bates et al. 2005, Onstad et al. 2018, Anderson et al. 2019). The extremely low number of pests that survive exposure to the toxins and/or pesticide are then expected to mate with the much higher number of susceptible pests that were reared on the refuge crop. However, due to the different management strategies employed between different crops, it is important to know how they may affect each other in areas where both crops (dry beans and corn) are grown in close proximity. It is therefore necessary to understand how adult moths that developed on beans are interacting with adult moths that developed on corn to evaluate the potential for synergistic IRM. Should they be interacting and breeding, it may be possible to use dry beans and corn as mutual refuges to help reduce the development of resistance in areas where both crops are grown in proximity.

We hypothesized that in regions where corn and dry beans are grown both crops are significant sources of adults that contribute to the gene pool. To test this hypothesis, it was first necessary to capture adult moths from an area where both beans and corn are grown, followed by determining the host plant used for development in the larval state via stable carbon isotope analysis. This is possible due to the strong isotopic differences of the stable isotopes of carbon (**δ**^13^C) found between C3 and C4 plants (Adams et al. 2016, Hambäck et al. 2016, Hobson et al. 2018). Tissues lacking large amounts of cellular turnover can be used to determine if the insect developed on C3 (dry beans) or C4 (corn) photosynthesizing plants. Traditionally, wings are used for stable isotope analysis, however difficulty in packaging the samples for analysis and loss of sample material due to damage led us to investigate if heads are a suitable alternative for carbon isotope analysis to differentiate between adult moths that developed on C4 and C4 plants.

## Material and methods

### Sample Collection

Sample collection was performed in central Michigan throughout adult western bean cutworm flight season (mid-July through mid-August) during 2017 and 2018. Nine to twelve traps were used at three to six sites (Figure 1). At each site, traps were placed in between adjacent fields of corn and beans that had been rotated from the other crop the previous year. Trap sites were spaced between 2.8 and 31.4 km with individual traps at each site placed approximately 150 to 1,000 m apart to ensure that they were operating independently. Corn and dry bean abundance within a 5 km and 50 km radius surrounding the trap sites was determined using Cropscape (Han et al. 2014, Han et al. 2014, Han et al. 2012, Boryan et al. 2011, Boryan et al. 2014) (Table 1 and Table 2). Individual trap sites were used for the 5 km radius determination. For the 50 km radius determination, a centralized point was selected between all traps and used to determine the crop abundance.

**Table 1.** Percentage of land use devoted to corn, dry beans and other in a 5Km radius from the collection site. The top portion of the table is for the samples collected in 2017 at the six sample sites. The bottom portion of the table is for the samples collected in 2018 at the three collection sites.

| Sample Collection (2017) |  |  |  |  |  |  |  |  |  |  |
| --- | --- | --- | --- | --- | --- | --- | --- | --- | --- | --- |
| Collection Site Information |  |  | 2016 |  |  |  | 2017 |  |  |  |
| Latitude | Longitude | Site Number | Corn (%) | Dry Beans (%) | Ratio (Corn:Beans) | Other (%) | Corn (%) | Dry Beans (%) | Ratio (Corn:Beans) | Other (%) |
| 43.1772 | -85.1629 | 1 | 39.8 | 6.7 | 6:1 | 53.5 | 26.0 | 5.2 | 5:1 | 68.8 |
| 43.2626 | -85.1819 | 2 | 43.6 | 6.0 | 7:1 | 50.4 | 20.8 | 3.4 | 6:1 | 75.8 |
| 43.2940 | -85.1812 | 3 | 19.2 | 3.4 | 6:1 | 77.4 | 23.3 | 3.5 | 7:1 | 73.2 |
| 43.3568 | -85.3790 | 4 | 29.1 | 8.6 | 3:1 | 62.3 | 28.1 | 11.2 | 3:1 | 60.7 |
| 43.3600 | -85.4160 | 5 | 42.4 | 15.7 | 3:1 | 41.9 | 28.5 | 11.2 | 3:1 | 60.3 |
| 43.3602 | -85.4624 | 6 | 19.3 | 7.8 | 2:1 | 73.0 | 20.1 | 6.0 | 3:1 | 73.8 |
| Sample Collection (2018) |  |  |  |  |  |  |  |  |  |  |
| Collection Site Information |  |  | 2017 |  |  |  | 2018 |  |  |  |
| Latitude | Longitude | Site Number | Corn (%) | Dry Beans (%) | Ratio (Corn:Beans) | Other (%) | Corn (%) | Dry Beans (%) | Ratio (Corn:Beans) | Other (%) |
| 43.2041 | -85.1538 | 1 | 23.8 | 3.6 | 7:1 | 72.7 | 22.9 | 3.8 | 6:1 | 73.3 |
| 43.3083 | -85.3198 | 2 | 18.9 | 3.7 | 5:1 | 77.4 | 21.7 | 2.6 | 8:1 | 75.8 |
| 43.3224 | -85.4293 | 3 | 24.5 | 6.3 | 4:1 | 69.2 | 25.0 | 4.9 | 5:1 | 70.2 |

**Table 2.** Percentage of land devoted to corn, dry beans and other in a 50Km radius from a centralized point located between all trapping sites (GPS coordinates: 43.2760, −85.2639) from 2016 to 2018.

| Year | Corn (%) | Dry Beans (%) | Ratio (Corn:Beans) | Other (%) |
| --- | --- | --- | --- | --- |
| 2016 | 13.34 | 0.63 | 21:1 | 86.03 |
| 2017 | 13.97 | 0.74 | 19:1 | 85.29 |
| 2018 | 14.18 | 0.76 | 19:1 | 85.06 |
| Average | 13.83 | 0.71 | 19:1 | 85.46 |

**Figure 1.**
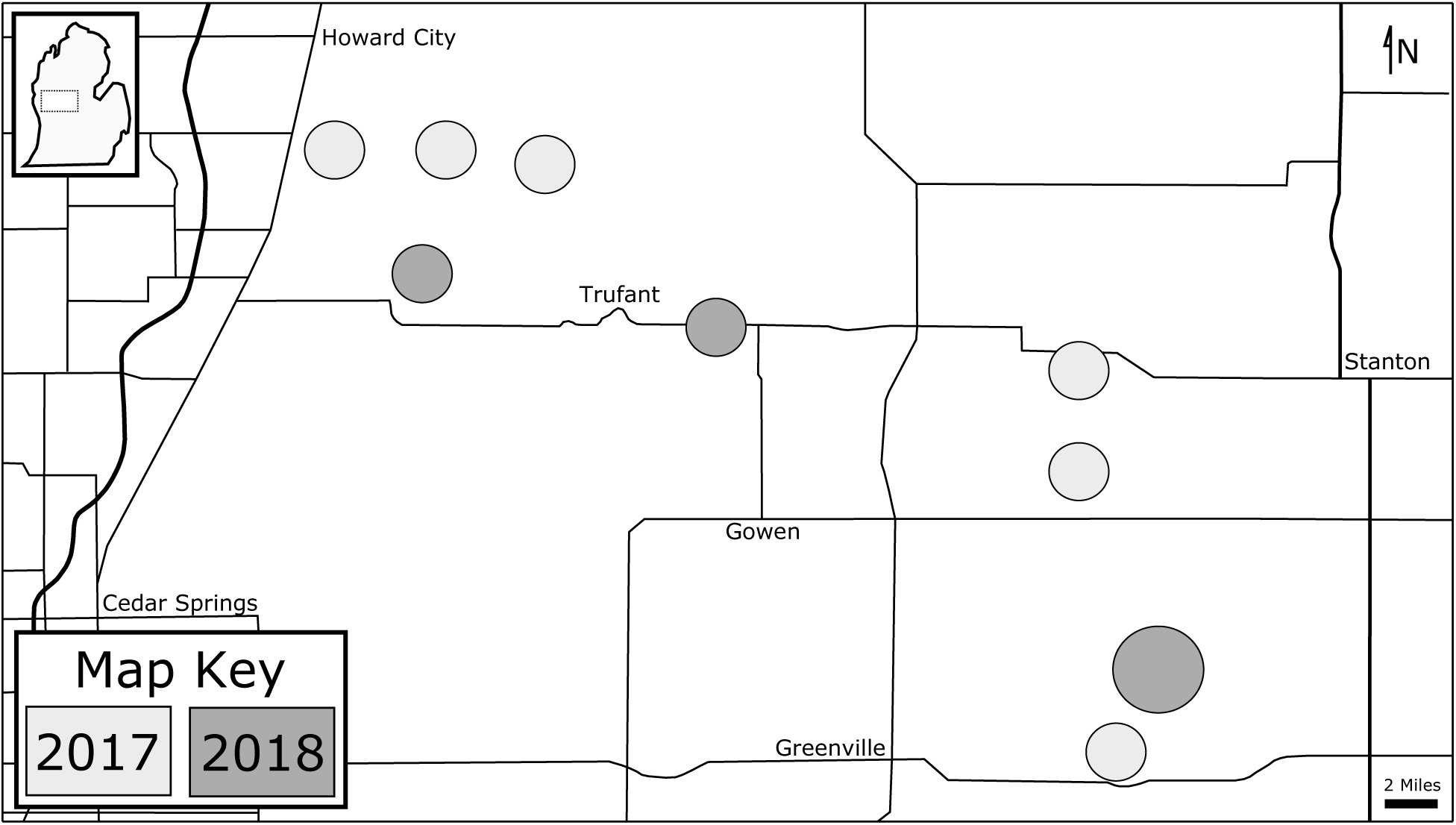
Map of capture sites in 2017 and 2018.

In 2017 and 2018, we used six “bucket” type universal moth traps (Great Lakes IPM Inc., Vestaburg, Michigan), that were baited with western bean cutworm sex pheromone lures (Scentry Biologicals Inc., Billings, Montana). In 2017, we unsuccessfully deployed six small portable LED blacklight traps (White et al. 2016) in an attempt to capture male and female moths. Therefore in 2018, three Quantum Black Light Traps (Leptraps LLC, Georgetown, Kentucky) were used in lieu of the portable LED blacklight traps. All traps contained a piece of insecticidal strip (Herocon Vaportape) to prevent moths from escaping or becoming damaged due to prolonged activity in the enclosed space. Traps were checked every morning and moths were placed in 50 mL centrifuge tubes containing 95% ethanol, returned to the lab and stored at −80°C until they were processed for isotope analysis.

### Stable Carbon Isotope Analysis

Samples were selected from periods of peak flight and given unique identifiers. Wings have historically been used for stable isotope analysis in moths due to slow carbon turnover (Adams et al. 2016). For this study we examined both wings and heads. Wings and heads were dried in an oven at 50°C until dry weight was constant (typically 48 hours) to evaporate any traces of ethanol and moisture. Dried wings, generally weighing between 1 to 4 mg, were ground and packaged into tin capsules for elemental and stable isotope analyses. Heads were packaged whole once dried.

Mass spectrometry was performed via standard protocols as described by Gonzalez-Meler et al. (2017). In short, samples were fully combusted to CO_2_ using a Costech (Valencia, CA) elemental analyzer equipped with a zero blank autosampler. Stable carbon isotope ratios were analyzed by an isotope ratio mass spectrometer operating in continuous flow (ThermoFinnigan Delta Plus XL equipped with Conflo III) connected to the elemental analyzer. Host plant carbon isotopic ratios were calibrated after host plant material was collected from the sample sites, dried at 60°C until constant dry weight, ground using tungsten microcontainers and a spex mill shaker, weighed into tin cups and analyzed as previously described. The stable carbon isotopic composition was expressed as a delta notation according to:

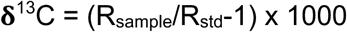

Where R is the ratio of ^13^C/^12^C of the sample and standard (std). Lab standards were calibrated against NIST standards using the Pee Bee Dolomite scale. Isotope analyses were performed at the University of Illinois at Chicago (USA) with a precision of 0.2 per mille (‰). The **δ**^13^C values from the plants, wings, and heads were then analyzed to determine individual larval hosts of adult moths. Samples that were tested for both head and wing tissues were then analyzed with Bartlett’s test of homogeneity of variances and paired t-tests.

## Results

A total of 3,264 adult moths were captured across the two years of trapping (Figures 2 and 3). In 2017, the portable LED light traps were successful in capturing multiple insect species, including other species of Lepidoptera. However, only 2 western bean cutworm moths were obtained whereas the pheromone traps captured 683 moths. In 2018, the Quantum blacklight traps captured 1,808 moths, and the pheromone traps captured 771 moths. In both 2017 and 2018, two peak periods of flight were observed with a clear peak occurring in mid-July and a second peak occurring in late July/ early August. Additionally, in 2017 and 2018, the use of pheromone traps resulted in only males being obtained and in 2018, through the use of blacklight traps, there were 1,431 males and 306 females collected with 71 having abdomens that were too damaged for sex to be determined. Overall 79% of all moths captured in 2018 via blacklight traps were male, 17% were female and 4% were undetermined.

**Figure 2.**
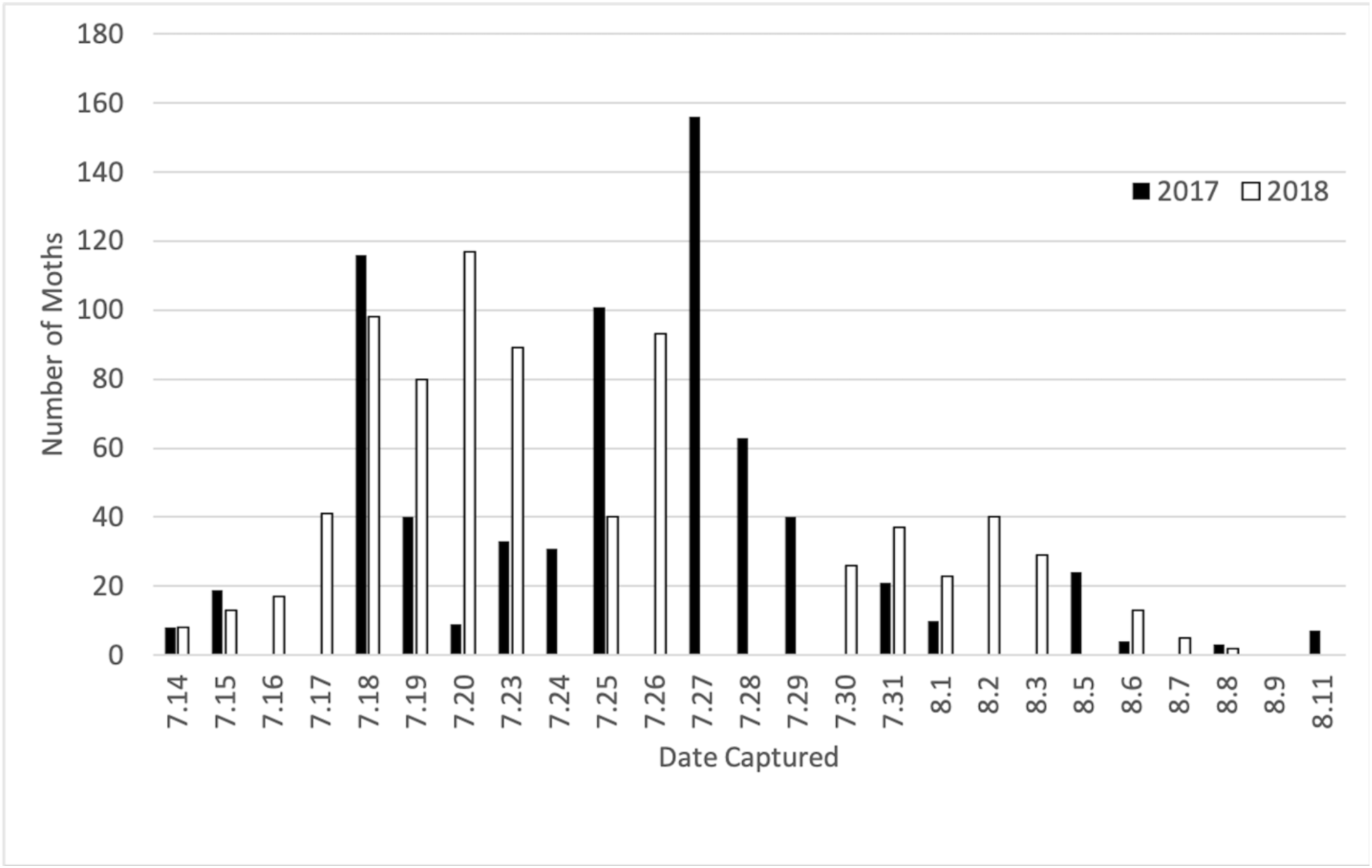
Number of male western bean cutworms collected with pheromone traps in 2017 (black) and 2018 (white).

**Figure 3.**
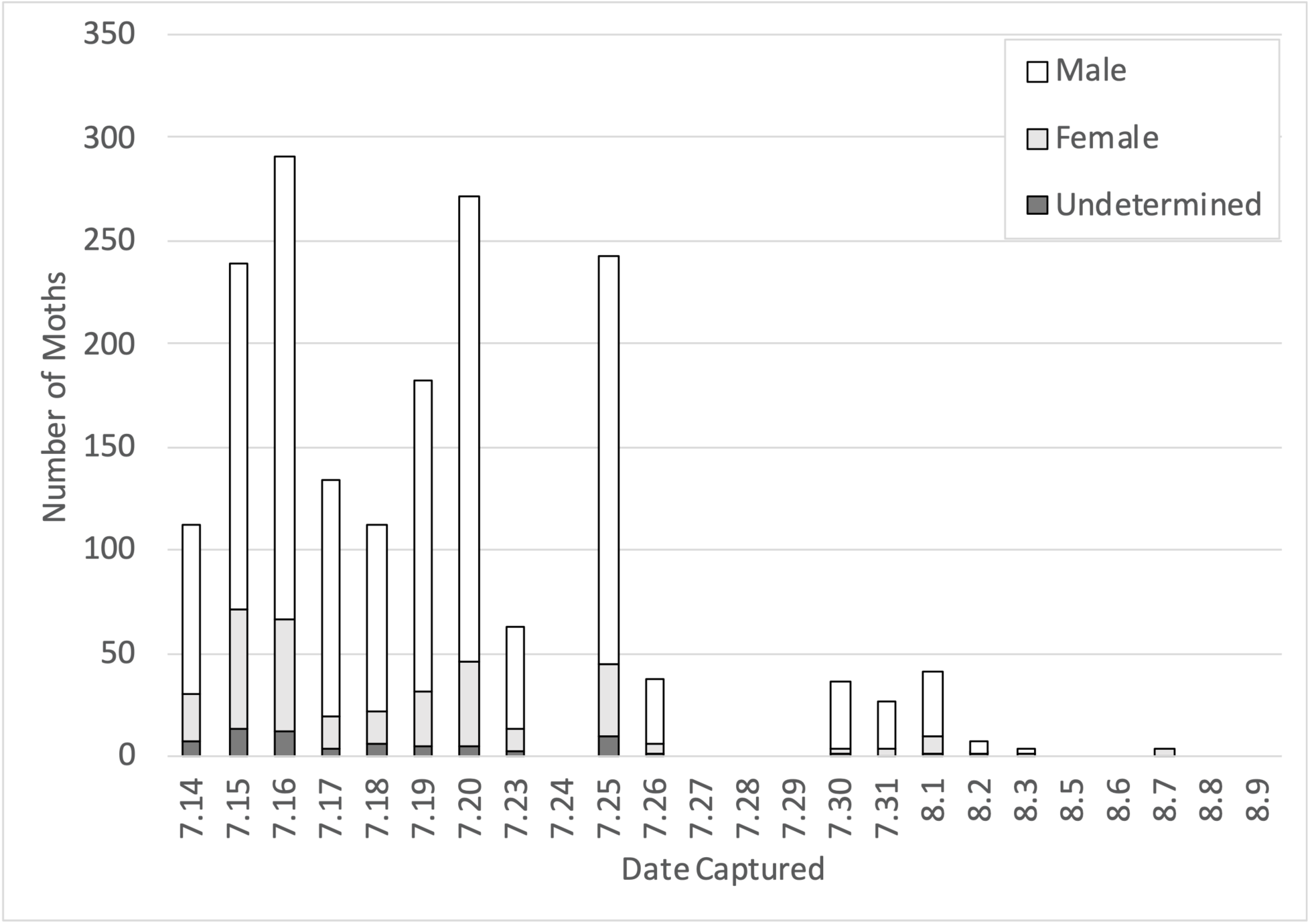
Number of western bean cutworms collected with Quantum blacklight traps in 2018.

Ten percent of the captured moths were used for stable carbon isotope analysis. A total of twenty-four moths were used to determine the effectiveness of head tissue compared to the more traditional wing tissue (Adams et al. 2016). Deviation between wings and heads for each sample was between 0.1 and 2.8 ‰ as shown in Figure 4. Overall, of the 349 moths that were tested, 346 (99.14%) of the moths presented a C4-like isotopic signature with a range of −11.0 to −15.1 ‰, whereas 3 (0.86%) of the moths presented a C3-like isotopic value with a range of −22.3 to −29.4 ‰. A slight upward bias (t-test: p = <10^−4^) was observed in the heads of moths presenting a C4-like isotopic signature, however a difference in the variance between the head and wing samples was not observed (Bartlett’s test: p = 0.47) (Figure 4). Statistical analyses were not performed on the individuals presenting a C3-like isotopic signature, because there were so few of them (*n* = 3).

**Figure 4.**
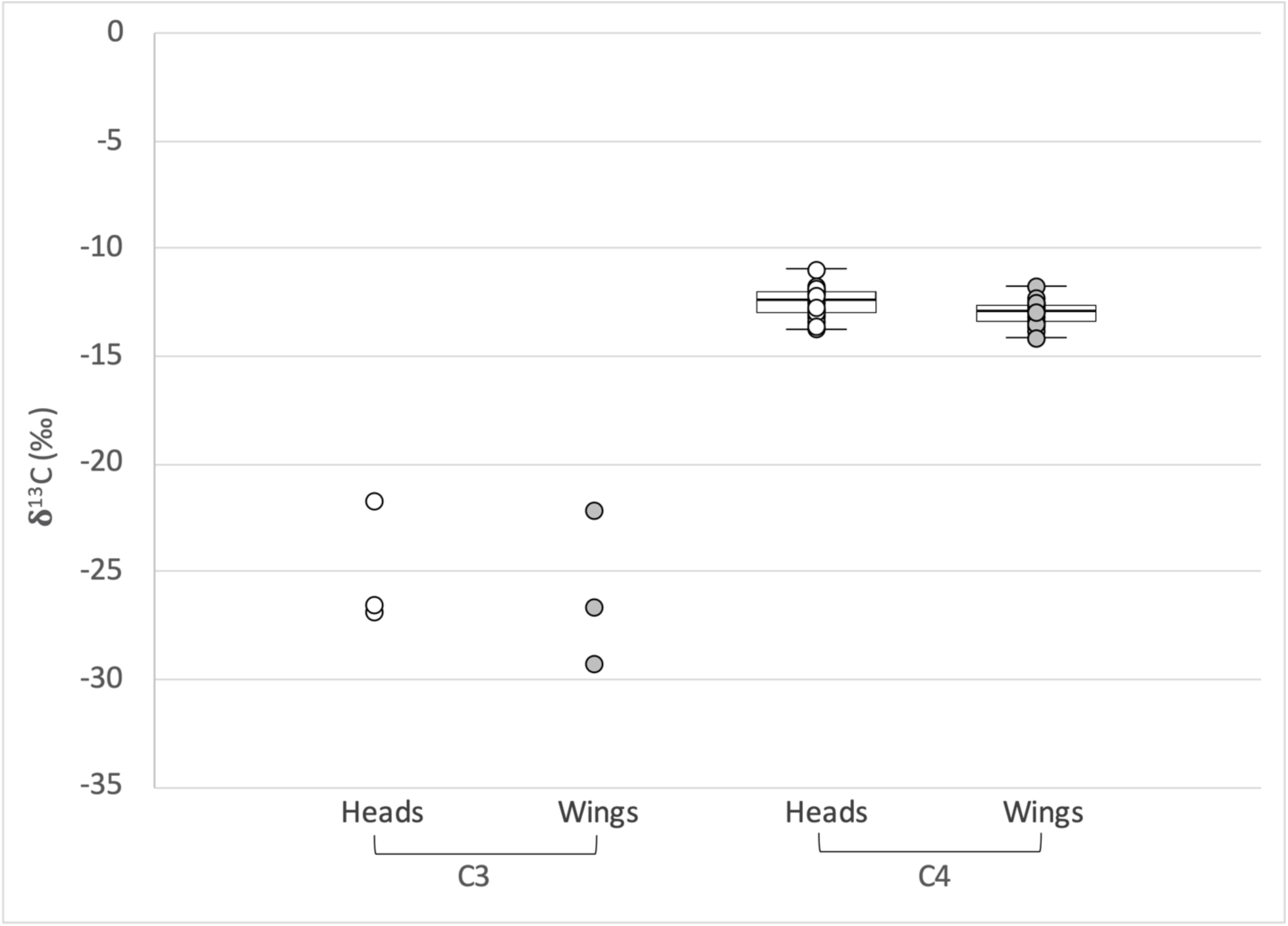
Depiction of the difference between **δ**^13^C (‰) values of heads and wings (* denotes a p-value of less than 0.05).

Crop distribution (Corn:Beans) within a 5 km radius surrounding trap sites for samples collected in 2017 ranged from 7:1 to 2:1 in 2016 and 7:1 to 3:1 in 2017. For samples collected in 2018 within 5 km of the trap sites, the ratios ranged from 7:1 to 4:1 in 2017 and 8:1 to 5:1 in 2018 (Table 1). When analyzing a 50 km radius from a centralized point between all traps, a ratio of 21:1 was observed in 2016, 19:1 in 2017, and 19:1 in 2018 (Table 2).

## Discussion

Through stable carbon isotope analysis, we found that less than 1% of the tested moths had isotopic signatures corresponding to C3 plants. This suggests that very few adult moths developed on dry beans as larvae. Additionally, we found evidence that the moths trapped in bean fields developed on C4 plants as larvae, demonstrating that they almost certainly developed on corn. These data suggest moths developed on corn as larvae are the primary contributors to the western bean cutworm gene pool in Michigan.

A possible explanation for this observation is that the large amount of corn grown in the area relative to dry beans creates a dilution reflected in the moth population. Subsequently, we analyzed the 5 km radii of the trap sites from 2016 to 2018 and observed that the ratios of corn to beans ranged from 8:1 to 2:1 (Table 1). If both crops were contributing equally to the adult population of western bean cutworm this would result in 11% to 33% of the moths having developed on dry beans as larvae, providing all captured moths developed within the immediate area. However, it was also possible that the moths developed further away from the trap sites.

We analyzed the 50 km radius surrounding the trap sites and found a ratio ranging from 21:1 to 19:1, or 4.5% to 5%, across all three years (Table 2). These values are still far greater than the <1% of C3 like moths observed. Therefore, while this dilution may play a role in the uneven ratio of moths that developed on corn as larvae compared to moths that developed on dry beans as larvae, it is unlikely that it is the only factor.

Another explanation for the low number of C3 like moths observed may be a preference for laying eggs on corn rather than dry beans. Previously, extension biologists have observed a preference for corn over dry beans when the corn has yet to tassel at the start of moth flight (Michigan State University Extension 2018). The stage of corn development varies widely based on planting time and factors such as precipitation and temperature and may have been affected by the abnormal growing conditions observed from 2016 to 2018 (Andresen 2017, Andresen 2018). While it is possible that the stage of corn development played an important role for the years we studied, this explanation fails to account for the consistent damage observed to dry beans on an annual basis (Smith et al. 2019).

A third explanation may be that larvae have a low success rate developing and overwintering under dry bean cultivation due to various unknown environmental factors. For example, soil type has been suggested to affect prepupae development and survivability in previous laboratory studies and foliar insecticides tend to be used more frequently in dry beans than in corn (Montezano et al. 2019, Difonzo et al. 2015). While we are unable to provide a precise reason for our observed results, they still suggest that corn fields act as an efficient supply of adult moths for the western bean cutworm, maintaining successful mating populations in the region and perpetuating the damage on dry beans the subsequent growing season.

Additionally, despite the presence of two flight peaks, it is unlikely that moths that developed on different host plants are flying at a separate time during the season as we sampled thoroughly from both peaks and found almost no moths that developed on C3 hosts. This is further supported by consistent monitoring performed by extension biologists in the area showing no evidence of differences in developmental timing (Tollini 2018, Springborn 2019). While we are unable to provide a precise explanation for the two flight peaks observed, insect emergence and crop development may have been affected by abnormal temperatures and variable precipitation observed both years (Andresen 2017, Andresen 2018). Our results are consistent with laboratory studies, which showed decreased survival of bean feeding larvae before the pupal stage of development when compared to larvae fed on corn (Montezano et al. 2019). Combined with data presented in this study, these results suggest that the environmental factors experienced by the recently established western bean cutworm in Michigan may result in them being unable to fully develop on dry beans, in which case, dry beans may be acting as a regional sink host for this pest.

The portable LED light traps (White et al. 2016) captured only two western bean cutworms and could not be used for the collection of our study species throughout the 2017 flight season. Subsequently, in 2018 commercial light traps were used, and proved more effective at trapping western bean cutworm moths. It is unclear why these particular portable LED traps were ineffective for the western bean cutworm, but it seems likely that either the wavelength or intensity of light or both were unattractive to this particular species. Our trap results suggest that large scale trapping efforts for the western bean cutworm should avoid the use of this portable LED trap.

In 2018, moths captured in light traps showed a large number of males compared to females. This may be due to more prevalent flight of male moths compared to female moths, making them more readily caught. While it has not been studied in the western bean cutworm, these results are consistent with previously observed male-biased sex ratios for samples caught via light traps (Altermatt et al. 2009, McDermott and Mullens 2018).

Stable carbon isotope analysis of the captured samples showed no observed overlap between moths that developed on C3 plants compared to moths that developed on C4 plants, and it was determined that both wings and heads can be used to distinguish between moths that developed on dry beans (C3 plants) compared to moths that developed on corn (C4 plants) (Figure 4). When performing the study, it was observed that wings were more difficult to package for analysis due to issues caused by poor integrity of the wing tissue and subsequent loss of sample material once dried and ground. This resulted in a newly trained scientist requiring eight hours to package twenty samples. When heads were used, however, they required much less handling time, and twenty samples only required two hours to package. Potentially, heads which experience more tissue turnover than wings, might show increased variation in isotope content due to feeding by adults. No difference in variance was observed between the **δ**^13^C values of heads and wing tissues in moths with C4-like isotopic signatures. While a slight upward bias of the **δ**^13^C values of heads and wing tissues in C4-like moths was observed it did not impact C3/C4 determination (Figure 4). Overall the use of head samples provided an adequate measurement for C3/C4 determination and allowed for a larger number of samples to be analyzed with less hands-on time required and increased productivity of the analysis by 75%.

Although there was an increased variance in the **δ**^13^C values of moths that developed on C3 plants compared to moths that developed on C4 plants, the values were within the expected range. Unlike C4 plants that perform photosynthesis across two different cell types to conserve water, C3 photosynthesis takes place in a single cell and is prone to various levels of gas exchange through the stomata depending on humidity and temperature. Subsequently, the variance of the **δ**^13^C in moths that developed on C3 plants is likely due to the relationship between the stomatal conductance and the individual plants photosynthetic rate (Farquhar et al. 1989). The small sample size may also be a factor affecting the variance of the **δ**^13^C values of moths that developed on C3 plants. Overall, our results would suggest that the determination of a C3/C4 host plant used for larval development via stable carbon isotope analysis in Lepidoptera could be facilitated through the use of head tissue, eliminating the issues caused by poor integrity of the wing tissue and subsequent loss of sample material, providing a more efficient way to perform population structure and migration studies and sampling in the future. However, further studies with different species and larger sample sizes are required to ensure that this is not a trait that is unique to the western bean cutworm. Additional studies are also required to ensure that head tissue can similarly be used for nitrogen and hydrogen isotopes with minimal effect due to adult feeding.

Our study suggests that various environmental factors experienced in central Michigan result in moths that developed on corn as larvae being the primary contributors to the mating pool and subsequently the next generation of western bean cutworm in this region. While currently undetermined, these environmental factors may include insecticide application and crop management such as acreage discrepancies, planting date, hybrid type, watering, etc. Additionally, soil type and nutrition as well as travel and migratory patterns of the adult moths may have an effect. Our results demonstrate that beans and corn cannot be used as mutual refuge in IRM in central Michigan and that further research is needed to determine proper IRM for areas where corn and dry beans are grown in close proximity.

## Acknowledgements

We thank two anonymous reviewers for their constructive comments on an earlier version of the manuscript. This work was supported in part by funds from the College of Science, Illinois Institute of Technology.

## Notes

### Competing Interest Statement

The authors have declared no competing interest.

### Summary of Updates

1. Data on the relative abundance of corn and beans in the local area. 2. Expanded discussion to consider alternative explanations for results.

